# Cytoneme-mediated signaling essential for tumorigenesis

**DOI:** 10.1101/446542

**Authors:** Sol Fereres, Ryo Hatori, Makiko Hatori, Thomas B. Kornberg

## Abstract

Communication between neoplastic cells and cells of their microenvironment is critical to cancer progression. To investigate the role of cytoneme-mediated signaling as a mechanism for distributing growth factor signaling proteins between tumor and tumor-associated cells, we analyzed EGFR and RET Drosophila tumor models. We tested several genetic loss-of-function conditions that impair cytoneme-mediated signaling. *diaphanous, Neuroglian, SCAR, capricious* are genes that cytonemes require during normal development. Genetic inhibition of cytonemes restored apical basal polarity to tumor cells, reduced tumor growth, and increased organism survival. These findings suggest that cytonemes traffic the signaling proteins that move between tumor and stromal cells, and that cytoneme-mediated signaling is required for tumor growth and malignancy.

**Summary:** Essential cytonemes for paracrine signaling in Drosophila tumors

## INTRODUCTION

Human tumors include transformed tumor cells, blood vessels, immune response cells, and stromal cells that together with the extracellular matrix (ECM) constitute a “tumor microenvironment” (Hanahan and Weinberg, 2000). The tumor microenvironment is essential for oncogenesis, cell survival, tumor progression, invasion and metastasis (Pietras and Östman, 2010; Roswall et al., 2018), and its stromal cells produce key drivers of tumorigenesis. Known drivers are growth factors (e.g. HGF, FGF, EGF, IGF-1, TGF-⃠ and Wnts), cytokines (e.g. IL-6, SDF) and pro-angiogenic factors (e.g. VEGF). These proteins may function as either autocrine, juxtacrine, or paracrine signals, but it is not known how they move into or within the tumor microenvironment.

Studies of tumor models in Drosophila exploit the experimental attributes of the fly that provide uniquely powerful ways to investigate tumorigenesis (Sonoshita and Cagan, 2017). We tested two models for the roles of cytonemes. Cytonemes are specialized, actin-based filopodia that extend between cells that produce and secrete signaling proteins and cells that receive them. The signaling proteins move along cytonemes and exchange at transient synapses that form where cytonemes contact target cells. These synapses are similar to neuronal synapses in both constitution and properties (Kornberg and Roy, 2014; Roy et al., 2014), and are necessary for paracrine FGF, BMP/Dpp, Hedgehog, Wnt/Wingless (Wnt/Wg), and Notch signaling during normal development of Drosophila epithelial tissues (González-Méndez et al., 2017; Huang and Kornberg, 2016; Roy et al., 2014).

EGFR activating mutations are drivers of several types of human cancers (Sibilia et al., 2007). Although elevated EGFR expression of wild type EGFR is not sufficient for tumorigenesis, additional genetic changes are necessary, such as over-expression of Perlecan, a heparan sulfate proteoglycan (HSPG) component of the ECM (Jiang and Couchman, 2003). In Drosophila, ectopic over-expression of Perlecan and EGFR in epithelial cells of the wing imaginal disc drives tumorigenesis (Herranz et al., 2014). Growth and metastasis of the epithelial cells requires crosstalk with closely associated mesenchymal myoblasts, which also proliferate abnormally when Perlecan and EGFR are over-expressed in epithelial neighbors. The crosstalk includes BMP/Dpp signaling in the epithelial cells and Wnt/Wg signaling in the mesenchymal cells (Herranz et al., 2014).

The RET gene is the primary oncogenic driver for MEN2 (multiple endocrine neoplasia type 2) syndrome. MEN2 is characterized by several types of neoplastic transformations, including an aggressive thyroid cancer called medullary thyroid carcinoma (MTC). Fly models that overexpress RET^MEN2^ phenocopy aspects of the aberrant signaling in MEN2-related tumors, such as activation of the SRC signal transduction pathway, which promotes migration and metastasis of tumorigenic cells. Screens for small molecule suppressors of the Drosophila RET^MEN2^ over-expression tumors identified compounds that are more effective than the drugs that are currently used for patients (Dar et al., 2012; Read et al., 2005).

In the work presented here we examined the role of cytoneme-mediated signaling in the EGFR-Pcn and the RET^MEN2^ models. Genetic inhibition of cytonemes by downregulation of *diaphanous, Neuroglian* and *SCAR*, genes previously shown to be essential in cytoneme-mediated signaling, reduced tumor growth and Dpp signaling, and suppressed lethality by 60% in the EGFR-Pcn tumor and by 30% in the RET^MEN2^ tumor. These results suggest that cytoneme-mediated signaling is necessary for tumor growth and that interfering with cytoneme-meditaed tumor-stromal cell signaling might be a therapy for tumor suppression.

## RESULTS

### Tumor cells and stromal cells extend cytonemes

Most of the wing imaginal disc is a columnar epithelium that generates both the adult fly wing and cuticle of the dorsal thorax. The disc also includes myoblasts that grow and spread over much of the dorsal basal surface of the columnar epithelium; these mesenchymal cells will generate the flight muscles of the adult. Tracheal branches also adjoin the basal surface of the columnar epithelium, and one branch, the transverse connective, sprouts a bud (the air sac primordium (ASP), Fig. 1A) that grows during the third instar period dependent on Dpp and FGF signaling proteins that are produced by the wing disc (Roy et al., 2014). The myoblasts relay Wg and Notch signaling between the disc and ASP (Huang and Kornberg, 2015). Cytonemes mediate and are essential for the Dpp, FGF, Wg, and Notch signaling during development (Kornberg, 2014).

**Figure 1.**
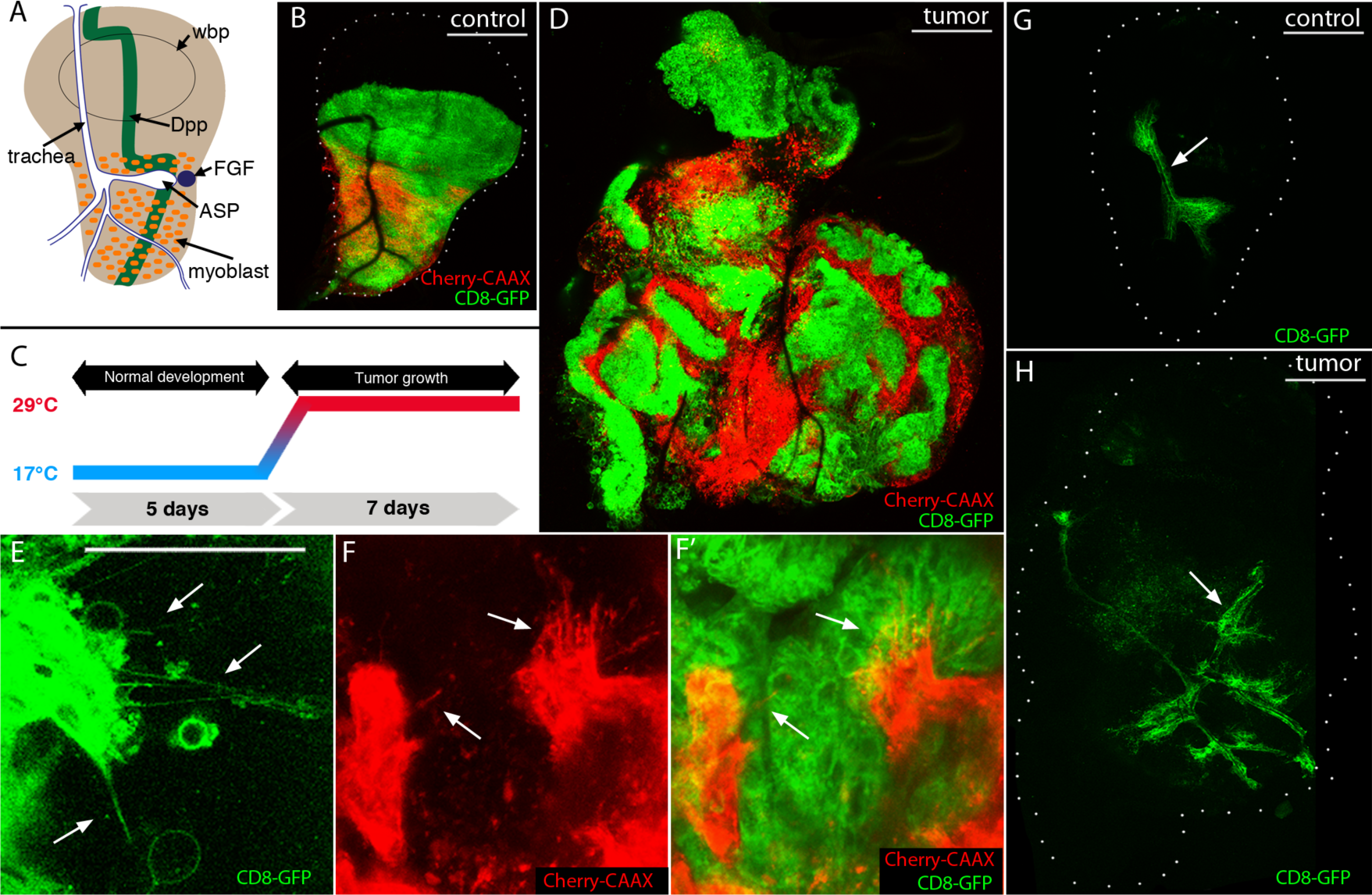
Cytonemes in EGFR-Pcn_i_ tumor and tumor-associated cells. (A) Cartoon of a 3^rd^ instar larval wing disc with wing blade primordia (wbp), disc-associated myoblasts (orange), trachea (white, outlined in blue), air sac primordium (ASP), Dpp expressing cells (green stripe), FGF expressing cells (blue circle). (B) Control wing disc expressing CD8:GFP in dorsal driven epithelial cells (*ap-Gal4*; green), and mCherry:CAAX in myoblasts ((*15B03-lexA*; red). (C) Schematic representation of tumor induction: animals developed for five days at 18°C with Gal4 repressed by Gal80, were transferred to 29°C to induce Gal4 and tumor growth for seven days (unless indicated otherwise). (D-F) Unfixed EGFR-Pcn tumor model wing discs. (D) Wing disc with tumor cells (CD8:GFP; green) and myoblasts (mCherry:CAAX; red). Genotype: *ap-Gal4,UAS-psq*_*RNAi*_*/115B03-lexA,lexO-Cherry-CAAX;UAS-EGFR,tub-Gal80*^*ts*^*/UAS-CD8:GFP*. Scale bars: 100 μm. (E) Cytonemes in the tumor epithelial cells (green, arrows); genotype: *ap-Gal4,UAS-psq*_*RNAi*_*/+;UAS-EGFR,tub-Gal80*^*ts*^*/UAS-CD8:GFP*. (F-F’) Cytonemes in myoblasts (red, arrows) extend towards epithelial cells (green); genotype: *ap-Gal4,UAS-psq*_*RNAi*_*/115B03-lexA,lexO-Cherry-CAAX;UAS-EGFR,tub-Gal80*^*ts*^*/UAS-CD8:GFP*; scale bars: 50 μm. (G-H) Unfixed wing discs with marked tracheal cells (green, arrows); (G) control, genotype: *btl-LHG, lexO-CD2- GFP*; (H) EGFR-Pcn tumor, genotype: *ap-Gal4,UAS-psq*_*RNAi*_*/btl-LHG,lexO-CD2-GFP;UAS-EGFR,tub-Gal80*^*ts*^*/+*. Excessive tracheal growth and ectopic branches indicated by arrows; scale bars: 100 μm.

To investigate whether cytonemes are also essential in tumorigenesis, we tested a cancer model that requires tumor-stroma interactions in which neoplastic transformation is driven by interactions between the wing disc epithelial cells and myoblasts (Herranz et al., 2014). Overexpression of wild type EGFR and Perlecan (*pcn*, a secreted heparan sulfate proteoglycan) in the columnar epithelium drives proliferation of the epithelial cells (which are genetically modified) as well as their genetically wild type myoblast neighbors. Tumorigenesis depends on Dpp signaling from the epithelial cells to the myoblasts. We first investigated if cytonemes are present in EGFR-Pcn overexpressing tumor cells. We induced the EGFR-Pcn tumor model (with *ap-Gal4,* an epithelial cell-specific driver) together with CD8:GFP, a membrane-tethered GFP protein (Figure 1C,D), and independently expressed membrane-tethered mCherry in the myoblasts (with *1151-lexA lexO-mCherry-CAAX*; a myoblast-specific driver). In this system, the epithelial cell membranes are marked with GFP fluorescence and the myoblast membranes are marked with mCherry fluorescence. We observed that, as previously reported (Herranz et al., 2014), the EGFR-Pcn tumor induces overgrowth and proliferation, producing multilayered masses of disorganized disc epithelial cells and myoblasts ((Herranz et al., 2014), Fig. 1D). Higher magnification imaging detected both epithelial cell and myoblast cytonemes, some of which appear to extend between the tumor and mesenchymal populations (Fig. 1E-F’). These results show that tumor cells and tumor-associated cells extend cytonemes and support the idea that cytonemes may facilitate signaling between these cell populations.

To determine if the tracheal system was also affected by the tumor conditions, we induced the EGFR-Pcn tumor and labelled the tracheal cells with membrane tethered GFP (with *LHG lexO-CD2:GFP*, a tracheal-specific driver (Shiga et al., 1996)). In the EGFR-Pcn tumor discs, the trachea were morphologically abnormal, and were more extensive and more branched than normal (Fig. 1G,H), suggesting that signaling to the genetically unmodified tracheal cells was also affected in the tumor discs.

### Dpp localizes to the tumor epithelial cell cytonemes

In normal development, Dpp produced by wing disc cells at the anterior/posterior compartment border is transported by cytonemes to target cells in both the wing disc and other tissues, and cytoneme deficits lead to developmental defects (Roy et al., 2014). In the EGFR-Pcn tumor model, Dpp signals from the mutant epithelial cells to drive myoblast expansion (Herranz et al., 2014). Dpp expression is upregulated in the epithelial cells (Fig. 2A,A’) and pMAD, the phosphorylated form of the Dpp signal transducer MAD, is highly enriched in the myoblasts (Fig. 2A”, 2A’”). This indication of Dpp signal transduction in the myoblasts is consistent with previous results showing that Dpp signaling in this stromal compartment is required for tumor growth (Herranz et al., 2014).

**Figure 2.**
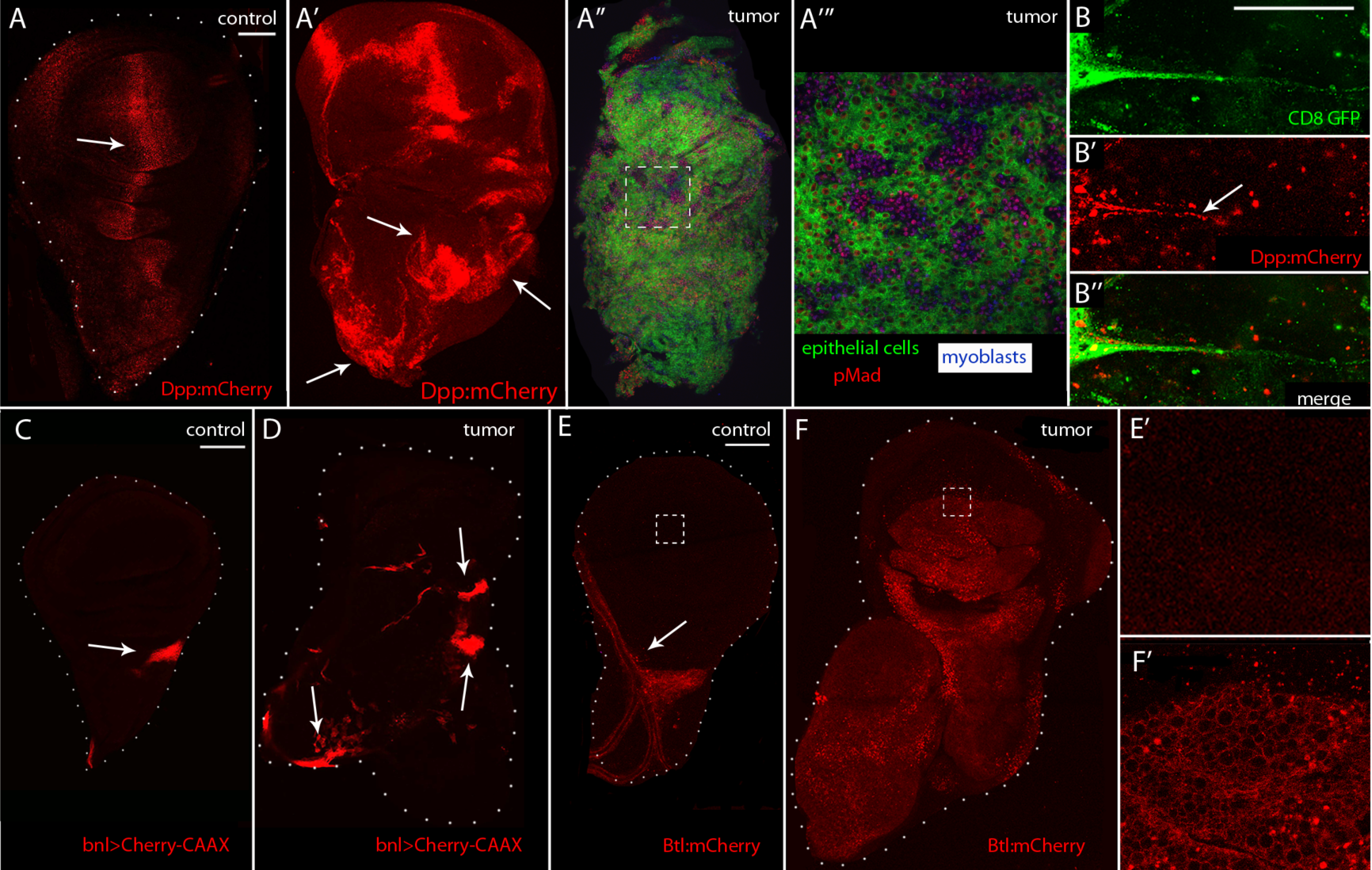
Dpp and FGF signaling pathways in the EGFR-Pcn tumor model. (A-B) Unfixed wing discs showing the Dpp distribution. (A) Control disc with Dpp (red, arrow). Genotype: *Dpp:mCherry*. Scale bar: 100⃠m. (A’) Disc with EGFR-Pcn tumor induced for 5 days expressing Dpp (red) in tumor cells. Genotype: *ap-Gal4,UAS-psq*_*RNAi*_*/Dpp:mCherry;UAS-EGFR,tub-Gal80*^*ts*^*/UAS-CD8:GFP*. Arrows indicate Dpp up-regulation. (A’’-A’’’) EGFR-Pcn wing showing epithelial tumor cells (green) fixed and stained with *α*-phosphorylated MAD (pMad, red) antibody to monitor Dpp signaling and *α*-Cut (blue) to label myoblasts. Genotype: *ap-Gal4,UAS-psq*_*RNAi*_*/+;UAS-EGFR,tub-Gal80*^*ts*^*/UAS-CD8:GFP* (A’’’) Higher magnification image of the box area in (A’’). (B-B’’) Cytoneme (green) extending from epithelial tumor cell (green) with Dpp:mCherry (red); arrow indicates Dpp:mCherry in cytoneme. Genotype: *ap-Gal4,UAS-psq*_*RNAi*_*/Dpp:mCherry;UAS-EGFR,tub-Gal80*^*ts*^*/UAS-CD8:GFP*. Scale bar: 50⃠m. (C-D) Unfixed wing discs with FGF/Bnl-expressing cells marked with Cherry-CAAX (red, arrows). (C) Control, genotype: *bnl-lexA,lexO-mCherry:CAAX/+*. (D) EGFR-Pcn tumor, genotype: *ap-Gal4,UAS-psq*_*RNAi*_*/+;UAS-EGFR,tub-Gal80*^*ts*^*/bnl-lexA,lexO-mCherry:CAAX*. Arrows indicate FGF up-regulation (red). (E-F’) Unfixed wing discs with FGFR/Btl distribution marked by Btl:mCherry (red, arrows). (E) Control, genotype: *Btl:mCherry*/+. FGFR/Btl is only expressed in the tracheal cells. (E’) Higher magnification image of the boxed area in (E). (F) 5 day EGFR-Pcn tumor disc, genotype: *ap-Gal4,UAS-psq*_*RNAi*_*/UAS-CD4-mIFP;UAS-EGFR,tub-Gal80*^*ts*^*/Btl:mCherry*. FGFR/Btl expression is upregulated in the tumor cells. (F’) Higher magnification image of the box area in (F). Scale bars: 100 μm.

To investigate how Dpp moves in the EGFR-Pcn model, we used CRISPR mutagenesis to tag the endogenous *dpp* gene with mCherry. We induced the EGFR-Pcn tumor in the Dpp:mCherry flies, and labeled the EGFR-Pcn tumor cells with CD8:GFP. Dpp:mCherry fluorescence was also present in the cytonemes of the epithelial tumor cells (Figure B,B’,B”), consistent with the idea that Dpp signaling is mediated by cytonemes in the EGFR-Pcn tumor model.

### FGF signaling between epithelial cells and myoblasts

Genetic ablation of myoblasts in the EGFR-Pcn tumor model reverts the tumor phenotype, suggesting that signaling from the myoblasts to transformed epithelial cells is required for tumor progression (Herranz et al., 2014). The signal has not been identified. Because the morphology of disc-associated tracheal branches is dependent on and sensitive to signals produced by the disc (Guha et al., 2009; Sato and Kornberg, 2002), and because growth of the tracheal branches increases in the EGFR-Pcn tumor model (Figure 1H), we investigated if FGF signaling is upregulated in tumor discs. Branchless (Bnl/FGF), a Dosophila FGF, is produced by a small, discrete group of disc cells (Fig 1A). Disc cells do not express the FGFR Breathless (Btl). In contrast, tracheal cells express Btl/FGFR, but not Bnl/FGF (Sato and Kornberg, 2002). To monitor FGF signaling in the EGFR-Pcn tumor model, we examined a Bnl/FGF reporter that expresses mCherry:CAAX in Bnl/FGF-expressing cells (Du et al., 2017). The number and location of Bnl/FGF-expressing cells increased in tumor discs (Fig. 2C,D). We also examined fluorescence of Btl:mCherry (a CRISPR-generated knock-in). Btl/FGFR is expressed by tracheal cells and has not been detected in larval wing disc epithelial cells ((Sato and Kornberg, 2002); Fig. 2E). In the tumor discs, however, Btl:mCherry fluorescence was present in the epithelial cells of the tumor (Fig. 2F). These results suggest that the tumor induces ectopic expression of Btl/FGFR and that excessive growth of the tracheal branches in this tumor model might be due to activation of the FGF signaling pathway.

To investigate the role of Bnl/FGF signaling in EGFR-Pcn tumorigenesis, we overexpressed a dominant negative FGFR mutant in the tumor cells to block FGF signaling (*UAS-Btl*^*DN*^). We compared disc morphology, Dpp signaling (monitored by anti-pMad antibody staining), and myoblast distribution (monitored by anti-Cut antibody staining, a specific mark of myoblasts (Blochlinger et al., 1993)) in control discs, EGFR-Pcn tumor discs, and EGFR-Pcn discs expressing Btl^DN^. Tumor discs were misshapen and six times larger than wild type discs, the number of Cut-expressing cells increased by four times, and anti-pMad staining was as well widespread and unpatterned (Fig. 3A,B,I). Expression of Btl^DN^ restored these features to normal. In discs over-expressing EGFR, Pcn, and Btl^DN^, morphology, the pattern of Dpp signaling, and distribution of myoblasts were indistinguishable from controls (Fig. 3C). These results indicate that inhibition of FGF signaling in the tumor cells suppressed both tumor growth and myoblast proliferation, and restored Dpp signal transduction to normal; they suggest that ectopic Btl/FGFR expression and Bnl/FGF signaling are key drivers of the tumor.

**Figure 3.**
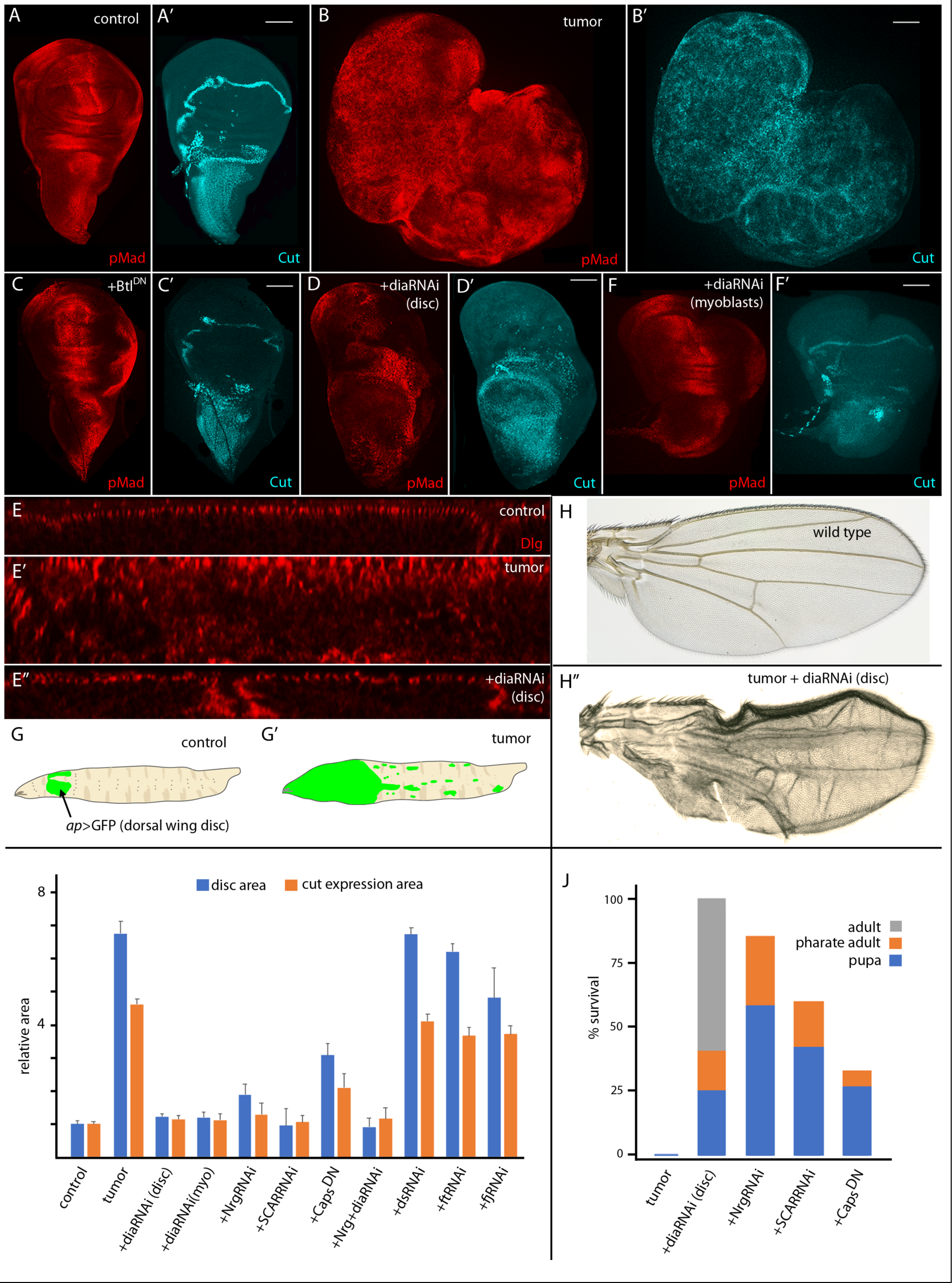
Tumor growth and signaling requires FGF signaling and *diaphanous*. (A-D) Fixed wing discs stained with ⃠-phosphorylated MAD (pMad, red) antibody to monitor Dpp signaling and ⃠-Cut (cyan) to label myoblasts. Scale bars: 100⃠m. (A,A’) Control. (B,B’) EGFR-Pcn tumor, genotype: *ap-Gal4,UAS-psq*_*RNAi*_*/+;UAS-EGFR,tub-Gal80*^*ts*^*/UAS-CD8:GFP*. (C,C’) EGFR-Pcn +Btl^DN^ expressed in epithelial cells. Genotype: *ap-Gal4,UAS-psq*_*RNAi*_*/+;UAS-EGFR,tub-Gal80*^*ts*^*/UAS-Btl*^*DN*^. (D,D’) EGFR-Pcn + diaRNAi in epithelial cells. Genotype: *ap-Gal4,UAS-psq*_*RNAi*_*/+;UAS-EGFR,tub-Gal80*^*ts*^*/UAS-dia*_*RNAi*_. (E-E”) Sagittal sections of fixed wing discs stained with ⃠-Dlg antibody (red) to mark the cell’s apical compartments. (E) Control. (E’) EGFR-Pcn tumor, genotype: *ap*-*Gal4,UAS-psq*_*RNAi*_*/UAS-CD8:GFP;UAS-EGFR,tub-Gal80*^*ts*^*/+*) (E’’) EGFR-Pcn + diaRNAi expressed in epithelial cells, genotype: *ap-Gal4,UAS-psq*_*RNAi*_*/+;UAS-EGFR,tub-Gal80*^*ts*^*/UAS-dia*_*RNAi*_. (F,F’) EGFR-Pcn expressed in epithelial cells + diaRNAi expressed in the myoblasts; fixed and stained with ⃠-pMad (red) and ⃠-Cut (cyan). Genotype: *ap-Gal4,UAS-psq*_*RNAi*_*/115B03-lexA;UAS-EGFR,tub-Gal80*^*ts*^*/ lexO-dia*_*RNAi*_. (G) Cartoon of a wild type larva depicting cells expressing GFP in the imaginal disc dorsal compartments (*ap-*Gal4; green, arrow). (G’) Cartoon of EGFR-Pcn tumor larva with overgrowth and metastasis throughout the larva (green). (H) Control adult wing. (H’) EGFR-Pcn + diaRNAi wing, genotype: *ap-Gal4,UAS-psq*_*RNAi*_*/UAS-CD8:GFP;UAS-EGFR,tub-Gal80*^*ts*^*/UAS-dia*_*RNAi*_. (I) Quantification of the total wing disc area (blue) and relative area of Cut-expressing cells (orange) of control, EGFR+Pcn tumor, tumor + diaRNAi expressed in epithelial cells, diaRNAi expressed in myoblasts, NrgRNAi, SCARRNAi, Caps^DN^, NrgRNAi+diaRNAi, dsRNAi, ftRNAi and fjRNAi larvae. Data was normalized to control. Student’s *t* test *P* values (*P*<0.05 to *P* >0.0005 for all except no significant difference for tumor + dsRNAi, ftRNAi and fjRNAi); *n* = 3-5 discs for each genotype. (J) Survival of EGFR-Pcn tumor, tumor + diaRNAi, NrgRNAi, SCARRNAi and Caps^DN^ larvae to pupal (blue), pharate adult (orange) and adult stage (gray). Student’s *t* test *P* values: (between *P*<0.05 and *P* >1.10^-8^) with *n* = 15-30 larvae for each genotype.

### Genetic suppression of tumor phenotypes

To assess the role of cytonemes in tumorigenesis, we examined discs in which cytonemes are impaired. Downregulation of *diaphanous* (*dia*), *Neuroglian* (*Nrg*), *SCAR* or *Capricious* (*Caps*) reduces the number and length of cytonemes, and reduces signaling in tracheal cells, myoblasts and wing disc cells (Bischoff et al., 2013; Huang and Kornberg, 2015; Roy et al., 2014). We expressed RNAi constructs to lower Dia, Nrg, SCAR and Caps levels in tumor cells, using conditions that represent partial loss-of-function and do not perturb cell polarity, cell viability, or cell cycle during normal development (Chen et al., 2017; Huang and Kornberg, 2015, 2016; Roy et al., 2014). We monitored wing discs for morphology, Cut expression, and pMAD. In contrast to discs with EGFR-Pcn tumors, discs with tumor cells that expressed diaRNAi in addition to EGFR and Pcn were morphologically less distorted, only 1.2 times larger than wild type, and the number and distribution of Cut-expressing cells was close to normal (Fig 3D,I). We also analyzed the apical-basal organization of the disc cells by motoring the distribution of Discs large (Dlg), which associates with the septate junction and localizes specifically to the apical compartment of the columnar epithelial cells. Sagittal optical sections of discs stained with anti-Dlg antibody reveal the specific apical distribution of Dlg characteristic of wild type cells is disorganized in EGFR-Pcn tumor discs (Fig. 3E,E’). Expression of diaRNAi in the tumor cells restored the Dlg distribution to normal (Fig. 3E”).

The presence of cytonemes in both the tumor (columnar epithelial) and mesenchymal (myoblast) cells, and the essential role of the myoblasts for tumor progression raises the possibility that myoblast cytonemes might also play an essential role in tumorigenesis. To investigate the role of myoblast cytonemes, we expressed diaRNAi (with *1151-lexA lexO-diaRNAi*) in the myoblasts of discs that overexpress EGFR and Pcn in the columnar epithelial cells. The morphology, Dpp signaling pattern and myoblast growth characteristic of the EGFR-Pcn tumors were suppressed (Fig. 3F,F’,I). This result is consistent with the idea that the myoblasts signal to the epithelial tumor cells and that this signaling is mediated by cytonemes. Although EGFR and Pcn expression in the EGFR-Pcn model (driven by *ap*-Gal4) is restricted to the dorsal compartment of the wing disc, the tumors grow extensively and metastasize (Fig. 3G,G”), and with no maturation beyond the larval stage (Herranz et al., 2014). The tumorous condition is 100% lethal. However, the conditions of diaRNAi expression that reduce cytonemes and cytoneme-mediated signaling suppressed lethality: all the EGFR-Pcn tumor-bearing larvae pupated, approximately 75% reached the pharate adult stage, and approximately 60% survived to adult stage (Fig. 3J). These surviving adults were fertile, and wing blade morphological defects were the only visible phenotype (Fig. 3H,H’). These results are consistent with the idea that cytoneme-mediated signaling is necessary for tumor growth and that interfering with tumor-stromal cell signaling suppresses many if not all aspects of tumorigenesis.

Expression of NrgRNAi, SCARRNAi, CAPS^DN^ in the epithelial cells of the EGFR-Pcn model reduced tumor growth, pMAD expression and number of Cut-expressing cells (Fig. 3I, Fig. 4A-D). Although these conditions improved survival, they were less effective than Btl^DN^ (Fig. 2) or diaRNAi (Figs. 3, 4). To test whether the ameliorative effects of RNAi constructs might be additive, we expressed NrgRNAi and diaRNAi simultaneously in EGFR-Pcn tumor cells. Expression of both NrgRNAi and diaRNAi decreased tumor growth (Fig. 3I), but compared to wild type wing discs, the discs were smaller and morphologically abnormal (Fig. 3I and 4), and no pupae developed to adult stage. We surmise that these effects might be due to the combined effects of insufficient function of the *Nrg* and *dia* genes. We also tested the roles of three genes that are essential for planar cell polarity (Sharma and McNeill, 2013): the *dachsous* (*ds*) and *fat* (*ft)* genes that encode cadherin family proteins, and *four-jointed* (*fj*) that encodes a transmembrane kinase. Expression of dsRNAi, ftRNAi or fjRNAi does not perturb cytoneme-mediated signaling (Huang and Kornberg, 2016), and expression of these RNAi lines in the tumor cells had no apparent effect on tumorigenesis (Fig. 4E-G).

**Figure 4.**
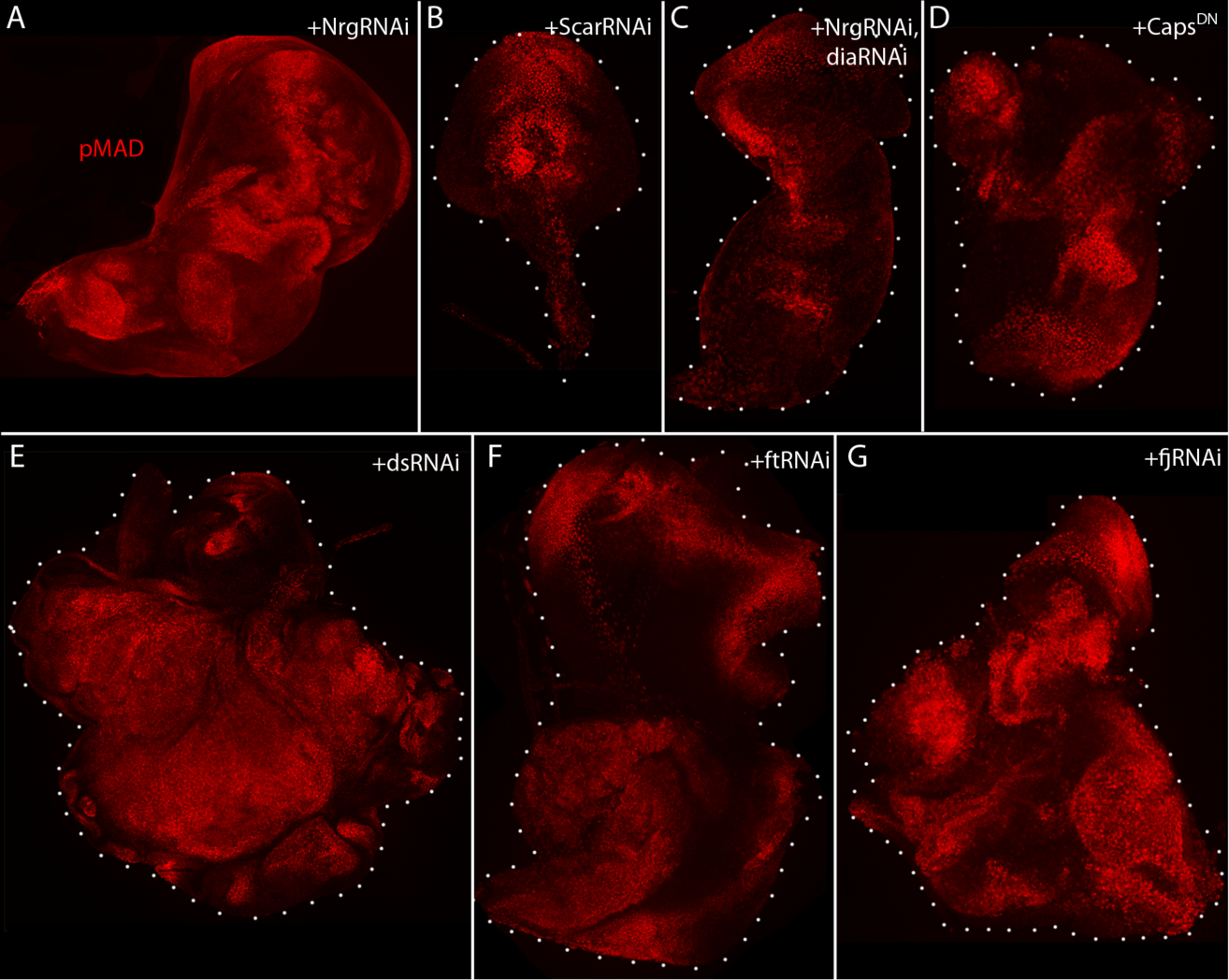
Conditions that ablate cytonemes decrease signaling, reduce tumor growth and promote survival. (A-D) Fixed wing discs stained with ⃠-pMad (red) antibody to monitor Dpp signaling. (A) Tumor + NrgRNAi, genotype: *ap-Gal4,UAS-psq*_*RNAi*_*/UAS-Nrg*_*RNAi*_*;UAS-EGFR,tub-Gal80*^*ts*^*/+*. (B) Tumor + SCARRNAi, genotype:*ap-Gal4,UAS-psq*_*RNAi*_*/+;UAS-EGFR,tub-Gal80*^*ts*^*/UAS-SCAR*_*RNAi*_. (C) Tumor + NrgRNAi, diaRNAi, genotype: *ap-Gal4,UAS-psq*_*RNAi*_*/UAS-Nrg*_*RNAi*_*;UAS-EGFR,tub-Gal80*^*ts*^*/UAS-dia*_*RNAi*_. (D) Tumor + Caps^DN^, genotype:*ap-Gal4,UAS-psq*_*RNAi*_*/UAS-CAPS*^*DN*^*;UAS-EGFR,tub-Gal80*^*ts*^*/+*. (E) Tumor + dsRNAi, genotype: *ap-Gal4,UAS-psq*^*RNAi*^*/+;UAS-EGFR,tub-Gal80*^*ts*^*/UAS-ds*_*RNAi*_‥ (F) Tumor + ftRNAi, genotype: *ap-Gal4,UAS-psq*^*RNAi*^*/+;UAS-EGFR,tub-Gal80*^*ts*^*/UAS-ft*_*RNAi*_. (G) Tumor + fjRNAi, genotype: *ap-Gal4,UAS-psq*^*RNAi*^*/+;UAS-EGFR,tub-Gal80*^*ts*^*/UAS-fj*_*RNAi*_

### Cytonemes in a RET-MEN2 tumor model

We investigated the role of cytonemes in the Drosophila RET-MEN2 tumor model developed by the Cagan lab (Dar et al., 2012). This model mimics the mis-regulation of signaling pathways that have been implicated in MEN2-related tumors. Overexpression of RET^MEN2^ in a discrete set of wing disc epithelial cells (with *ptc-Gal4*) resulted in >4X increase in the number of *ptc-*expressing cells and a 9X increase in the portion of the disc that consists of *ptc-*expressing cells (Figure 5A-C) (Dar et al., 2012). Approximately one-half of the animals survive to the pupal stage, but none survived to adult. We tested whether diaRNAi expression in the RET-mutant cells affects tumor growth and survival, and observed that excessive growth of the *ptc-*expressing cells was suppressed by more than 4X (Fig. 5C,D). Approximately two-thirds of the animals developed to the pupal stage, and one-third survived to adult. These flies were wild type in appearance, without any apparent visible abnormalities. These results are consistent with a general role for cytonemes in tumorigenesis and tumor progression.

**Figure 5.**
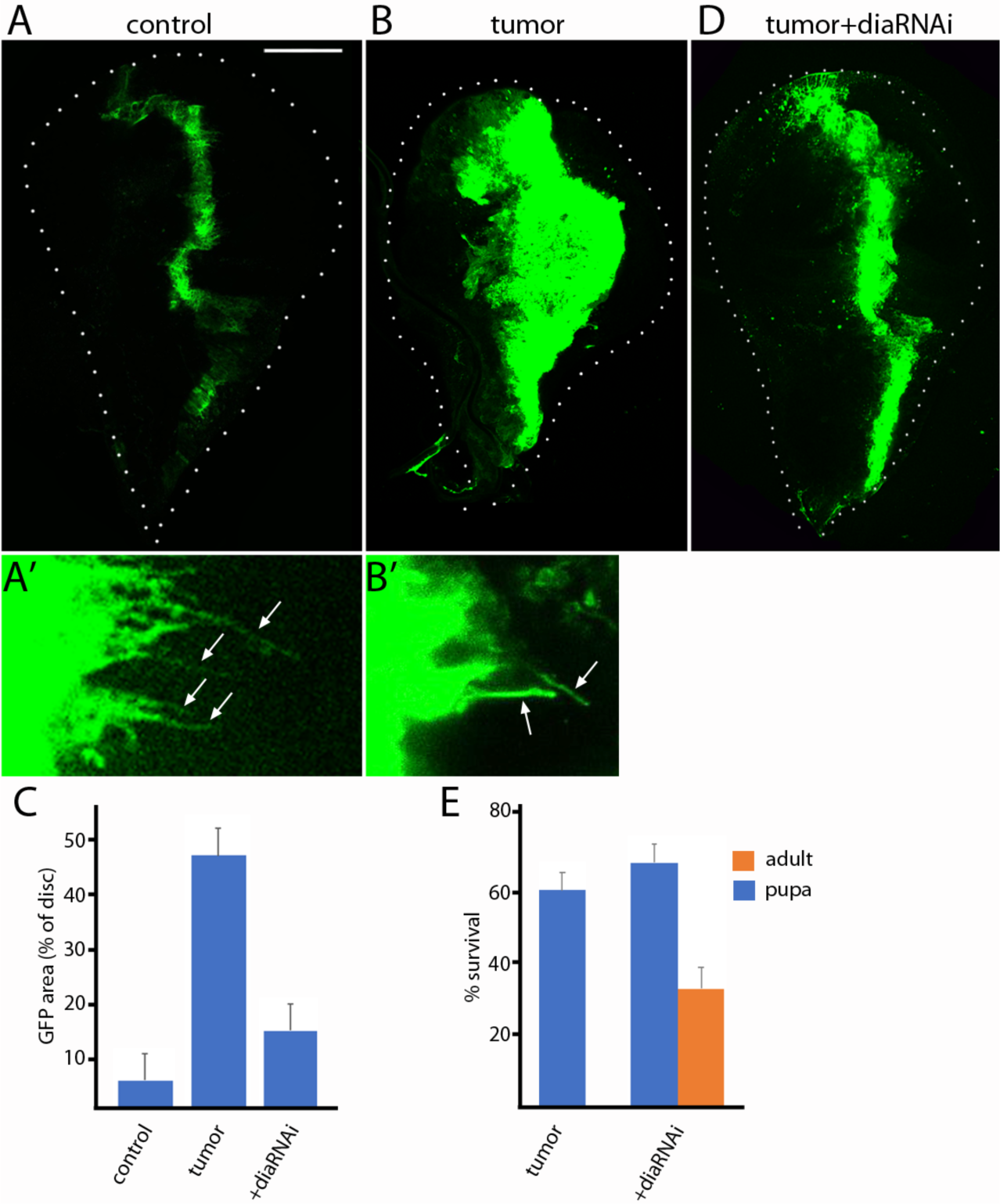
RET-MEN2 tumor growth and survival depends on cytonemes. (A-B’) Unfixed wing discs expressing CD8:GFP (green) driven by *ptc-Gal4.* Scale bar: 100⃠m. (A) Control. (A’) Cytonemes in wild type cells (green, arrows). (B) RET tumor, genotype: *ptc-Gal4,CD8:GFP;UAS-RET*^*MEN2*^*/UAS-CD8:GFP.* (B’) Cytonemes in RET tumor cells (green, arrows). (C) Quantification of the area of the disc expressing GFP (% of disc) in control, RET tumor and RET + diaRNAi discs. Significance was analyzed using Student’s *t*-test (*P*<1.10^-5^) and a total *n* of 4 discs. (D) Unfixed RET + diaRNAi wing disc expressing CD8:GFP (green). Genotype: *ptc-Gal4,CD8:GFP;UAS-RET*^*MEN2*^*/UAS-dia*_*RNAi*_. (E) Survival of RET-tumor and RET+ diaRNAi larvae to pupae (blue) and adults (orange). Significance using Student’s *t*-test (*P*<0.05 for RET+ diaRNAi pupae and *P*<0.005 for RET+ diaRNAi adults) with *n* = 30-40 larvae for each genotype.

## DISCUSSION

The tumor microenvironment is a niche that responds to signaling proteins produced by tumor cells and supplies growth factors that support tumor growth and metastasis (Quail and Joyce, 2013). Much ongoing work seeks inhibitors of tumorigenesis that target the signaling molecules and growth factors, their signal transduction pathways, and the stromal cells of the microenvironment (Heinemann et al., 2014; Presta et al., 2017). The target we investigated is the mechanism that transfers the signaling molecules and growth factors between tumor cells and the stromal cells of the niche.

Our previous work established that during Drosophila development, paracrine signaling by the signaling proteins/growth factors Dpp, FGF, Wg, Notch, Hedgehog, and Notch is mediated by cytonemes. Cytonemes are specialized filopodia that extend between signal producing and signal receiving cells, making synaptic contacts where the signaling proteins transfer from producing to receiving cells. To extend this work to tumorigenesis, we applied the strategies and tools we developed for our previous studies to ask if cytonemes are present in the tumor microenvironment, and if genetic conditions that inhibit cytoneme function and cytoneme-dependent signaling in normal development also inhibit tumorigenesis.

In a EFGR-Pcn tumor model, we found that cytonemes extend from both Drosophila tumor and stromal cells (Figs. 1 and 2). This is consistent with previous studies that reported increased signaling between tumor and stromal cells in this model (Herranz et al., 2014), and with the presence of cytonemes in many other contexts of paracrine signaling (Bischoff et al., 2013; Chen et al., 2017; Eom et al., 2015; Inaba et al., 2015; Roy et al., 2014; Sanders et al., 2013; Stanganello et al., 2015). We confirmed that Dpp is expressed by the tumor cells (Fig. 2; (Herranz et al., 2014)), and found that ectopic FGF signaling also has an essential role in this tumor (Figs. 2 and 3). The implied functional connections between the EGF, Dpp, and FGF signaling pathways have not been identified.

We also found conditions that impair cytonemes and cure flies of lethal tumors in both EGFR-Pcn and RET models. We selected four genes from among the more than twenty that are known to be essential for cytoneme-mediated signaling (Bischoff et al., 2013; Huang and Kornberg, 2016; Roy et al., 2014). *dia, Nrg, SCAR*, and *Caps* are recessive lethal genes whose functions can be partially reduced in genetic mosaics without affecting viability, cell shape, or the cell cycle, but are necessary for cytoneme function. Downregulating any one of these genes improved viability in the EGFR-Pcn model; *dia* inhibition was most effective in both models, and the cures that they effected suggest that cytoneme-mediated signaling, which might be a general mechanism for tumorigenesis in a variety of cancers, might also be a potential target for therapy.

The high degree of evolutionary conservation with human proteins make Drosophila a clinically relevant platform for understanding mechanisms human disease, and Drosophila tumor models have successfully identified new therapeutic candidates for colorectal, lung and thyroid and stem-cells derived cancers (Bangi et al., 2016; Levine and Cagan, 2016; Markstein et al., 2014). Our work provides proof principle for tumor suppression by interfering with cytoneme-mediated signaling.

## Acknowledgements

We thank: Drs. S. Cohen, H. Herranz, S. Roy, and R. Cagan and Bloomington Stock Center for fly stocks and R. Cagan for his help and advice; all members of Kornberg lab for discussion and constructive suggestions. Funding: NIH grant CA205580 to T.B.K. Author contributions: conceptualization - SF, RH, TBK; investigation – SF, RH, MH; Writing - SF, TBK. Competing interests: none.

## Experimental Procedures

### Drosophila stocks and husbandry

Flies were reared on standard cornmeal and agar medium at 29°C, unless otherwise stated. *ap*- Gal4 UAS-*psq*_*RNAi*_/*CyO*; UAS-EGFR *tub*-Gal80^ts^ from S. Cohen (Herranz et al., 2014), UAS-RET^MEN2^ from R. Cagan (Dar et al., 2012), Btl:mCherry and Bnl-lexA, lexO-mCherry:CAAX from S. Roy (Du et al., 2017). lexO-dia_RNAi_ from H. Huang. *ptc*-Gal4, *15B03*-lexA, lexO-Cherry:CAAX, UAS-CD8:GFP, UAS-dia_RNAi_, UAS-Nrg_RNAi_, UAS-SCAR_RNAi_, UAS-Caps^DN^, UAS-ds_RNAi_, UAS-ft_RNAi_, UAS-fj_RNAi_ from Bloomington Stock Center.

### Dpp:mCherry

The Dpp:mCherry transgene has mCherry inserted C-terminal to Dpp amino acid 465 (Entchev et al., 2000), with Leu-Val linkers inserted before and after a mCherry coding sequence deleted of its stop codon. The transgene was generated by CRISPR mutagenesis as follows:

### Dpp:mCherry donor vector

Left homology arm fragment contains overlapping sequence with PBS-SK vector and mCherry. The mCherry fragment contains overlapping sequence with the left homology arm and right homology arm. The right homology arm fragment contains overlapping sequence with mCherry and PBS-SK vector. The three fragments were stitched together and cloned into PBS-SK vector using Gibson Assembly (NEB). The resulting vector is designated as Dpp:Cherry donor vector.

Left arm homology sequence was amplified from wild-type genomic DNA using: L-arm-fwd: cggtatcgataagcttgatcaccttgccgcacaaatacatatac L-arm-rev: CCTCGCCCTTGCTCACCATCTCCAGGCCACCGCCCTCTCCGGCAGACACGTCCCGA

The mCherry tag was amplified using: mCherry-fwd:TGTCTGCCGGAGAGGGCGGTGGCCTGGAGATGGTGAGCAAGGGCGAGGAGGATAAC Cherry-rev:CGCTTGTTCCGGCCGCCCTTCTCTAACTTGTACAGCTCGTCCATGCCGC

The right arm homology sequence was amplified from wild-type genomic DNA using: R-arm-fwd:GGACGAGCTGTACAAGTTAGAGAAGGGCGGCCGGAACAAGCGGCAGCCGA R-arm-rev:ccgggctgcaggaattcgatGTCATTATTCGGTTATGCTCTCGCTAG

### pCFD-3 gRNA vector

gRNA sequence: CGCTCCATTCGGGACGTGTCTGG

The gRNA sequence without the PAM was cloned into pCFD-3 vector obtained from Addgene.

### Dpp:Cherry CISPR lines

pCFD-3 gRNA vector and Dpp:mCherry donor vector were co-injected into Cas9 expressing flies (nanos-Cas9) by Rainbow Transgenics. The resulting CRISPR-generated flies were screened and verified by sequencing. The Dpp:mCherry fly is viable and homozygous and hemizygous normal.

### EGFR-Pcn tumor

EGFR-Pcn tumors were induced as described by Herranz et al, (Herranz et al., 2014) by overexpression of EGFR and down-regulation of *pipsqueak* (*psq*), which leads to increased levels of Pcn. Female flies from the stock *ap-Gal4,UAS-psq*_*RNAi*_*/CyO;UAS-EGFR,tub-Gal80*^*ts*^ were crossed to males of the corresponding genotypes at 18°C, and were cultured at 18°C to maintain Gal80 repression of Gal4 and allow normal development. After 5 days larvae were transferred to 29°C to induce Gal4 expression and tumor growth. Tumor growth was induced for 7 days, unless otherwise indicated, whereupon larvae were dissected for live imaging or immunostaining, or were maintained at 29°C for survival studies. To control for possible effects on Gal4 expression, all tested genotypes had three UAS transgenes – either UAS-EGFR, UAS-psqRNAi and UAS-CD8:GFP for tumor flies, or UAS-EGFR, UAS-psqRNAi and additional RNAi for comparisons.

### RET tumor model

Female flies from the RET^MEN2^ stock (Dar et al., 2012) were crossed at room temperature to either *ptc*-*Gal4*, 2xUAS-CD8:GFP males or *ptc*-*Gal4*, UAS-CD8:GFP, UAS-diaRNAi males. For analysis of discs, embryos from one day collections were transferred to 29°C and cultured to third instar stage. For survival comparisons, animals were cultured at 25°C.

### Live imaging of wing imaginal discs

Wing discs with trachea attached were dissected in cold phosphate-buffered saline (PBS), placed on a coverslip and mounted upside-down on a coverslip on a depression slide as described (Huang and Kornberg, 2015). Samples were imaged with a Leica TCS SPE confocal or an Olympus FV3000 inverted confocal laser scanning microscope.

### Immunohistochemistry

Wing discs were dissected in cold PBS and fixed in 4% formaldehyde for 20 minutes. After extensive washing, the samples were permeablized with PBST (PBS + 0.3% TritonX-100), blocked for 1h with PBST+3%BSA blocking buffer, and incubated with primary antibodies previously diluted in blocking buffer overnight at 4°C. The following primary antibodies were used: α-pMad (Abcam), α-Discs large (Dlg), α-Cut and α-⃠-galactosidase (Developmental Studies Hybridoma Bank). Secondary antibodies were conjugated to Alexa Fluor 405, 488, 555, or 647. Samples were mounted in Vectashield and imaged with a Leica TCS SPE confocal or an Olympus FV3000 inverted confocal laser scanning microscope.

## References

Bangi, E., Murgia, C., Teague, A.G.S., Sansom, O.J., and Cagan, R.L. (2016). Functional exploration of colorectal cancer genomes using Drosophila. Nat. Commun. 7.

Bischoff, M., Gradilla, A.C., Seijo, I., Andres, G., Rodriguez-Navas, C., Gonzalez-Mendez, L., and Guerrero, I. (2013). Cytonemes are required for the establishment of a normal Hedgehog morphogen gradient in Drosophila epithelia. Nat Cell Biol 15, 1269–1281.

Blochlinger, K., Jan, L.Y., and Jan, Y.N. (1993). Postembryonic patterns of expression of cut, a locus regulating sensory organ identity in Drosophila. Development 117, 441–450.

Chen, W., Huang, H., Hatori, R., and Kornberg, T.B. (2017). Essential basal cytonemes take up Hedgehog in the Drosophila wing imaginal disc. Development dev.149856.

Dar, A.C., Das, T.K., Shokat, K.M., and Cagan, R.L. (2012). Chemical genetic discovery of targets and anti-targets for cancer polypharmacology. Nature 486, 80–84.

Du, L., Zhou, A., Patel, A., Rao, M., Anderson, K., and Roy, S. (2017). Unique patterns of organization and migration of FGF-expressing cells during Drosophila morphogenesis. Dev. Biol. 427, 35–48.

Entchev, E. V, Schwabedissen, A., and González-Gaitán, M. (2000). Gradient formation of the TGF-beta homolog Dpp. Cell 103, 981–991.

Eom, D.S., Bain, E.J., Patterson, L.B., Grout, M.E., and Parichy, D.M. (2015). Long-distance communication by specialized cellular projections during pigment pattern development and evolution. Elife 4.

González-Méndez, L., Seijo-Barandiarán, I., and Guerrero, I. (2017). Cytoneme-mediated cell-cell contacts for hedgehog reception. Elife 6.

Guha, A., Lin, L., and Kornberg, T.B. (2009). Regulation of Drosophila matrix metalloprotease Mmp2 is essential for wing imaginal disc:trachea association and air sac tubulogenesis. Dev. Biol. 335, 317–326.

Hanahan, D., and Weinberg, R.A. (2000). The hallmarks of cancer. Cell 100, 57–70.

Heinemann, V., Reni, M., Ychou, M., Richel, D.J., Macarulla, T., and Ducreux, M. (2014). Tumour-stroma interactions in pancreatic ductal adenocarcinoma: Rationale and current evidence for new therapeutic strategies. Cancer Treat. Rev. 40, 118–128.

Herranz, H., Weng, R., and Cohen, S.M. (2014). Crosstalk between epithelial and mesenchymal tissues in tumorigenesis and imaginal disc development. Curr. Biol. 24, 1476–1484.

Huang, H., and Kornberg, T.B. (2015). Myoblast cytonemes mediate Wg signaling from the wing imaginal disc and Delta-Notch signaling to the air sac primordium. Elife 4, e06114.

Huang, H., and Kornberg, T.B. (2016). Cells must express components of the planar cell polarity system and extracellular matrix to support cytonemes. Elife 5.

Inaba, M., Buszczak, M., and Yamashita, Y.M. (2015). Nanotubes mediate niche-stem-cell signalling in the Drosophila testis. Nature 523, 329–332.

Jiang, X., and Couchman, J.R. (2003). Perlecan and Tumor Angiogenesis. J. Histochem. Cytochem. 51, 1393–1410.

Kornberg, T.B. (2014). Cytonemes and the dispersion of morphogens. Wiley Interdiscip. Rev. Dev. Biol. 3, 445–463.

Kornberg, T.B., and Roy, S. (2014). Communicating by touch--neurons are not alone. Trends Cell Biol 24, 370–376.

Levine, B.D., and Cagan, R.L. (2016). Drosophila Lung Cancer Models Identify Trametinib plus Statin as Candidate Therapeutic. Cell Rep.

Markstein, M., Dettorre, S., Cho, J., Neumuller, R.A., Craig-Muller, S., and Perrimon, N. (2014). Systematic screen of chemotherapeutics in Drosophila stem cell tumors. Proc. Natl. Acad. Sci. 111, 4530–4535.

Pietras, K., and Östman, A. (2010). Hallmarks of cancer: Interactions with the tumor stroma. Exp. Cell Res. 316, 1324–1331.

Presta, M., Chiodelli, P., Giacomini, A., Rusnati, M., and Ronca, R. (2017). Fibroblast growth factors (FGFs) in cancer: FGF traps as a new therapeutic approach. Pharmacol. Ther. 179, 171–187.

Quail, D., and Joyce, J. (2013). Microenvironmental regulation of tumor progression and metastasis. Nat. Med. 19, 1423–1437.

Read, R.D., Goodfellow, P.J., Mardis, E.R., Novak, N., Armstrong, J.R., and Cagan, R.L. (2005). A drosophila model of multiple endocrine neoplasia type 2. Genetics 171, 1057–1081.

Roswall, P., Bocci, M., Bartoschek, M., Li, H., Kristiansen, G., Jansson, S., Lehn, S., Sjölund, J., Reid, S., Larsson, C., et al. (2018). Microenvironmental control of breast cancer subtype elicited through paracrine platelet-derived growth factor-CC signaling. Nat. Med. 24, 463–473.

Roy, S., Huang, H., Liu, S., and Kornberg, T.B. (2014). Cytoneme-Mediated Contact-Dependent Transport of the Drosophila Decapentaplegic Signaling Protein. Science. 343, 1244624–1244624.

Sanders, T.A., Llagostera, E., and Barna, M. (2013). Specialized filopodia direct long-range transport of SHH during vertebrate tissue patterning. Nature 497, 628–632.

Sato, M., and Kornberg, T.B. (2002). FGF is an essential mitogen and chemoattractant for the air sacs of the Drosophila tracheal system. Dev. Cell 3, 195–207.

Sharma, P., and McNeill, H. (2013). Fat and Dachsous Cadherins. Prog. Mol. Biol. Transl. Sci. 116, 215–235.

Shiga, Y., Tanaka-Matakatsu, M., and Hayashi, S. (1996). A nuclear GFP/beta-galactosidase fusion protein as a marker for morphogenesis in living Drosophila. Dev. Growth Differ. 38, 99–106.

Sibilia, M., Kroismayr, R., Lichtenberger, B.M., Natarajan, A., Hecking, M., and Holcmann, M. (2007). The epidermal growth factor receptor: From development to tumorigenesis. Differentiation 75, 770–787.

Sonoshita, M., and Cagan, R.L. (2017). Modeling Human Cancers in Drosophila. In Current Topics in Developmental Biology, pp. 287–309.

Stanganello, E., Hagemann, A.I.H., Mattes, B., Sinner, C., Meyen, D., Weber, S., Schug, A., Raz, E., and Scholpp, S. (2015). Filopodia-based Wnt transport during vertebrate tissue patterning. Nat. Commun. 6, 5846.

